# Polyploidy enhances desiccation tolerance in the grass *Microchloa caffra*

**DOI:** 10.1101/2023.06.20.545583

**Authors:** Rose A. Marks, Paula Delgado, Givemore Munashe Makonya, Keren Cooper, Robert VanBuren, Jill M. Farrant

## Abstract

Desiccation tolerance evolved recurrently across diverse plant lineages to enable survival in water limited conditions. Many resurrection plants are polyploid and several groups have hypothesized that polyploidy enabled the evolution of desiccation tolerance. However, due to the vast evolutionary divergence between resurrection plant lineages, the rarity of desiccation tolerance, and the prevalence of polyploidy in plants, this hypothesis has been difficult to test. Here, we surveyed variation in morphological, reproductive, and desiccation tolerance traits across natural populations of a single species that has differing ploidies and tested for links between polyploidy and resilience. We sampled multiple populations of the resurrection grass *Microchloa caffra* across an environmental gradient ranging from mesic to xeric in South Africa. We describe two distinct ecotypes of *M. caffra* that occupy different ends of the environmental gradient and exhibit consistent differences in ploidy, morphological, reproductive, and desiccation tolerance traits in both field and common growth conditions. Interestingly, plants with more polyploid genomes were consistently more desiccation tolerant, less reproductive, and larger than plants with smaller genomes and lower ploidy. These data suggest that polyploidy enhances desiccation tolerance and that stronger selective pressures in increasingly xeric sites may play a role in maintaining and increasing desiccation tolerance.

## INTRODUCTION

Desiccation tolerant organisms can withstand nearly complete drying of their tissues without dying (Bewley, 1979; Bewley and Krochko, 1982; Oliver et al., 2020). Plants with this ability in their vegetative tissues are commonly referred to as resurrection plants because of their dramatic ability to revive from what appears to be a dead condition. Desiccation tolerance likely evolved in plants more than 500 mya and played a critical role in enabling the transition from aquatic to terrestrial environments by early land plants (Oliver et al., 2000). Throughout evolutionary time, desiccation tolerance was lost (or suppressed) in the vegetative tissues of most vascular plants, but was maintained in many early diverging lineages (e.g., bryophytes, ferns, fern allies) and specialized tissues (e.g., seeds, spores, and pollen) (Proctor and Tuba, 2002; Oliver et al., 2005). Desiccation tolerance then re-evolved convergently in the vegetative tissues of diverse vascular plants that occupy extremely arid habitats, likely through rewiring of ancestral pathways maintained in seeds and other specialized tissues (Costa et al., 2017; VanBuren, 2017; VanBuren et al., 2017). As a result, extant resurrection plants are phylogenetically and morphologically diverse, with representatives in every major land plant lineage except gymnosperms. Resurrection plants are found on all seven continents (Alpert, 2006) and often occur in closely intertwined communities on isolated rock outcrops (Porembski, 2011) and other extremely arid habitats. Resurrection plants have received growing research attention over the past two decades (Tebele et al., 2021) due to their astonishing resilience, but many questions remain regarding the evolutionary history, genetic mechanisms, and natural diversity of desiccation tolerance (Marks et al., 2021).

Plant evolution has been continually shaped by polyploidy and genome duplication, and an estimated 70% of flowering plants have experienced at least one whole-genome duplication event in their evolutionary history (Soltis et al., 2014). This chromosomal redundancy may function to enhance adaptability to extreme environmental conditions, as the duplicated genes offer a broader genetic basis for the evolution of potential stress tolerance mechanisms (Comai, 2005). Polyploidy can also foster the evolution of novel traits by providing raw genetic material for gene diversification and subfunctionalization, which in turn, can lead to neofunctionalization and the development of new and emergent phenotypes (Wendel, 2000). As a result, polyploidy plays a crucial role in driving plant diversification, speciation, and adaptation to varied ecological niches (Otto, 2007; Te Beest et al., 2012; Soltis et al., 2015). Many resurrection plants are polyploid and several groups have hypothesized that polyploidy enabled the evolution of desiccation tolerance (Giarola et al., 2017) because genome duplication could allow seed related regulatory networks to neofunctionalize, enabling tolerance in vegetative tissues.

However, testing this hypothesis is challenging because of the vast evolutionary divergence between resurrection plants, the rarity of tolerant lineages, the prevalence of polyploidy across plants, and the association of polyploidy with numerous other adaptive traits. A natural system of variable ploidies within a single species would minimize confounding factors and could be used to test for associations between polyploidy and desiccation tolerance.

The diverse and climate resilient Chloridoid subfamily of grasses has more than 40 desiccation tolerant species (Marks et al., 2021), and an estimated 90% of species within this clade are polyploid (Roodt and Spies, 2003). Despite the prevalence of polyploidy within this lineage, some desiccation tolerant Chloridoid grasses such as *Oropetium thomaeum, O. capense, Tripogon minimus, T. spicatus*, and *T. loliiformis* are diploid with very small genomes of ∼220-270 Mb (Mehra and Sharma, 1975; Bartels and Mattar, 2002; VanBuren et al., 2015). In contrast, other desiccation tolerant Chloridoid grasses including *Eragrostis nindensis* and *Sporobolus stapfianus* are tetraploid (Pardo et al., 2020; Chávez Montes et al., 2022) and there is no clear pattern between polyploidy and desiccation tolerance within the Chloridoid grasses. However, most studies on desiccation tolerance within this group have focused on a single accession or ecotype, and few studies have surveyed natural variation in desiccation tolerance across multiple populations of a single species.

Here, we surveyed natural variation in ploidy, desiccation tolerance, morphological, and reproductive traits in a single Chloridoid grass species. We sampled multiple populations and accessions of the resurrection grass *Microchloa caffra* across an environmental gradient spanning ∼500 km in the Mpumalanga and Limpopo provinces of northeastern South Africa. We employed both field and common garden studies to distinguish between plastic and genetic differences in phenotypes. Our results provide evidence of heritable differences in desiccation tolerance, morphology, and reproduction within a single species. Phenotypic variation in desiccation tolerance and morphology was positively associated with genome size and elevated ploidy, suggesting that gene and genome duplication may play a role in the evolution of desiccation tolerance. Differences in desiccation tolerance also parallel the environmental gradient, with plants from dryer habitats generally exhibiting higher desiccation tolerance than plants from wetter habitats. We describe multiple trait associations between desiccation tolerance and other functional traits that point towards relevant trade-offs and can guide applied objectives. These findings provide insight into how organisms acquire and enhance desiccation tolerance on short evolutionary timescales and push us one step closer to the ultimate translation of desiccation tolerance into sensitive crops species–a major motivation of research in the field.

## MATERIALS AND METHODS

### Study species

*Microchloa caffra Nees* is perennial grass in the Chloridoid subfamily of Poaceae commonly known as pincushion grass. *Microchloa caffra* is distributed across Southern Africa from Uganda to South Africa with the highest density of plant in northeastern South Africa (Figure 1). Plants occur in seasonally dry summer rainfall areas in shallow soils on rocky outcrops or inselbergs, locally referred to as ruwari. The vegetative tissues of *M. caffra* are desiccation tolerant and can survive repeated cycles of de- and re-hydration within a single growing season (personal observation RAM).

**Figure 1.**
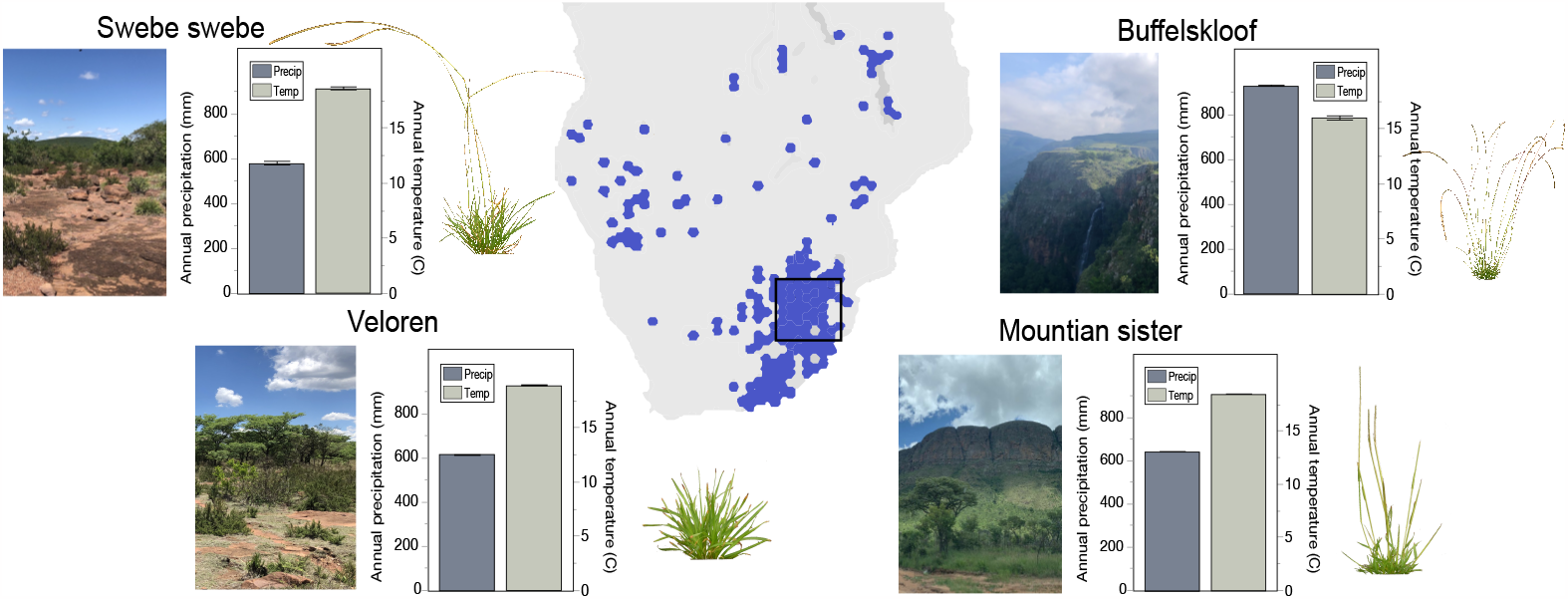
Distribution of *Microchloa caffra* taken from www.gbif.org. The study area spans ∼500 km in Mpumalanga and Limpopo provinces of northeastern South Africa. Representative images of the four sampling regions and plants are included, and bioclimatic data on annual precipitation and temperature are plotted from WorldClim (Fick and Hijmans, 2017). Multiple populations were sampled within each region.

### Field collections and phenotyping

Live specimens and seeds of *M. caffra* were collected from 11 populations in four major regions across Mpumalanga and Limpopo provinces of South Africa from Dec. 2021 to Jan. 2022 (Figure 1). Voucher specimens were deposited in the National Herbarium Pretoria (specimen number PRE1004810-0). The GPS coordinates and elevation of each population were logged with a Garmin 64csx GPS and bioclimatic data on annual precipitation and temperature were extracted from WorldClim (Fick and Hijmans, 2017) based on the GPS coordinates of each site using the R package “raster” (Table S1).

Seven plants from each population were phenotyped in the field, three of which were collected, potted, and brought into cultivation for seed bulking and further experimentation. To phenotype plants in the field, we measured the length and width of three randomly selected leaves and panicles from each plant and counted the total number of panicles on each plant. Measurements were taken to the nearest 0.5 mm.

### Seed bulking and germination tests

Seeds were collected from each of the 33 accessions (3 plants from 11 populations) and transported to Michigan State University under United States Department of Agriculture (USDA) permit #537-22-37-10071 and according to the specifications in a Material Transfer Agreement established between Drs. Jill M. Farrant, Robert VanBuren, and Rose A. Marks. Seeds were cold stratified for 2 weeks at 4°C and then germinated on a standard propagation mix of 50:50 redi-earth and suremix under controlled conditions in growth chambers set to ∼400 μmol light with a 16-hour photoperiod and 28/18°C day/night temperature. Depending on availability, 5 or 10 seeds were sown for each of the 33 accessions. Germination percentage was recorded at 4 weeks and seedlings were then transplanted into individual pots for experimental treatments.

Replicates of each accession were split into two sets, one for morphological phenotyping and the other for desiccation treatment.

### Phenotyping seed grown plants

Plants selected for phenotyping were photographed every two weeks for 12 weeks (beginning at 6 weeks old and extending to 18 weeks old) using a Nikon D5600 DSLR Camera with a 35 mm lens. Profile and overhead images were taken with scale and color standards included in each photograph. At 11 weeks old, plants were manually measured to quantify leaf and panicle length and width—paralleling field measurements. The number of panicles on each plant was manually counted when plants were 8 months old and plants were maintained for seed production. The time to panicle emergence and seed maturity were monitored. The first panicles emerged when plants were between 6 and 8 weeks old and the entire cycle from seed to seed was completed within ∼4-5 months, depending on the accession.

### Genome size and ploidy estimations

The genome size of each accession was estimated by flow cytometry to generate 2C DNA values. Healthy leaf tissue was harvested from each accession and submitted to Plantploidy.com where it was run on a BD Accuri™ C6 Plus Flow Cytometer. Hosta was used as an internal reference.

Ploidy was estimated for a single representative *M. caffra* accession with a haploid genome size of ∼1 Gb using a K-mer based approach (Ranallo-Benavidez et al., 2020). Approximately 34 Gb of PacBio high fidelity (HiFi) was generated for the reference accession from Buffelskloof. K-mers were counted from the high-quality circularized consensus reads using Jellyfish (V2.3.0) (Marçais and Kingsford, 2011) with default parameters and a K-mer length of 21 bp. A histogram of the K-mer distribution was used as input for GenomeScope2.0 and the monoploid genome size, ploidy, and likely origin of polyploid events (allo or auto) were calculated. A monoploid genome size of ∼300 Mb was used as a baseline to estimate ploidy of all the accessions.

### Screening desiccation tolerance phenotypes in field collected plants

Three *M. caffra* plants from each of the 11 populations (33 total) were collected from the field, potted, and transported to the University of Cape Town where they were maintained in a climate-controlled greenhouse for one month. Selected accessions from the two most environmentally extreme sites were targeted for desiccation assays. Six individual plants from Swebe Swebe and five from Buffelskloof were transferred to a growth room maintained at 20°C with a 16-hour photoperiod. After one week, they were subjected to a desiccation treatment and their ability to recover was assessed. Plants were watered to full soil saturation, after which water was withheld and the plants dehydrated naturally over the following 9 days. Plants were then rehydrated by rewatering with dH_2_O and their recovery was assessed.

Plants were sampled every 48 hours during the drying process and again 48 hours after rehydration. At each sampling timepoint, maximum quantum yield of Photosystem II (*F*_*v*_*/F*_*m*_) was measured, and leaf tissue was collected to quantify relative water content (RWC) and fixed for transmission electron microscopy (TEM). All measurements were taken in triplicate. Briefly, *F*_*v*_*/F*_*m*_ was measured on dark adapted leaves using a PAR-Fluorpen FP 110/D at 455 nm to quantify photosystem II efficiency. RWC was quantified by measuring leaf mass immediately after collection (fresh mass), after 48 hours submerged in water in darkness at 4°C (turgid mass), and again after 48 hours in a 70ºC drying oven (dry mass). RWC was calculated as (fresh mass - dry mass)/(turgid mass - dry mass). Samples were fixed for TEM and processed following (Cooper and Farrant, 2002). Briefly, leaves were fixed overnight at 4°C in 2.5% glutaraldehyde in 0.1 m phosphate buffer (pH 7.4) containing 0.5% caffeine. Next, samples were post-fixed by washing in 1% osmium phosphate buffer and then dehydrated in graded ethanol (30, 50, 70, 85, 95, and 100%) and washed twice in 100% acetone. The samples were incubated overnight at 4° in a 1:1 solution of acetone to Spurr’s resin. The concentration of Spurr’s resin was gradually increased over the course of 4 days until 100% Spurs resin was reached. The resulting samples were placed into molds and allowed to harden at 60°C for 24 h. Fixed tissues were sectioned using a Richart Ultracut S Ultramicrotome and mounted on copper grids. Sections were stained with uranyl acetate and lead citrate for contrast and visualized on an FEI Tecnai T20 transmission electron microscope.

### Screening desiccation tolerance phenotypes in seed grown plants

Comprehensive desiccation assays were performed on a replicated set of all accessions grown from seed under common conditions. Progeny from each of the 33 accessions were subjected to a desiccation treatment at Michigan State University similar to the treatment imposed on field collected plants at University of Cape Town. Six-week-old plants were watered to full soil saturation, after which water was withheld and plants were allowed to dry naturally and desiccate over the course of 4 weeks. Plants were sampled once a week on Tuesday at 14hr00. After 4 weeks, plants were rehydrated with dH_2_O and sampled 24, 48, 96, and 192 hours after rehydration. At each sampling timepoint, overhead and profile photographs were taken and *F*_*v*_*/F*_*m*_ was quantified on dark adapted leaves using a Opti-Sciences OS30p+ chlorophyll fluorometer with the default test parameters. RWC was measured immediately before rehydration (as described above) to confirm that plants had reached a desiccated state.

Overhead photographs from the final recovery timepoint (192 hours post rehydration) were analyzed using ImageJ to quantify the percentage of tissue that survived and recovered from desiccation. Images were segmented in ImageJ following the protocol described at https://edis.ifas.ufl.edu/publication/HS1382. Briefly, plant canopies were separated from the background using the color thresholding tool with hue, saturation, and brightness (HSB) set to specific values. We set the thresholds for hue to 1-170, saturation to 69-255, and brightness to 50-255. We cleared the background, removed outliers (points with a radius <10 pixels) and manually removed any remaining soil particles in the image using the eraser tool on three settings (300, 50, and 20 px). Next, senesced tissue was isolated from healthy tissue by setting the thresholding parameters to hue 1-35, saturation 0-255, and brightness 0-255. To isolate healthy recovered tissue, the thresholding parameters were set to hue 35-255, saturation 0-255, and brightness 0-255. The areas of the entire canopy, the senesced tissue, and the recovered tissue were calculated by converting the isolated areas into a binary image (black and white) and computing the area. Percent recovery for each replicate was calculated as the proportion of recovered tissue relative to the entire canopy.

### Statistical analyses

All statistical analyses were run in JMP 15 (SAS Institute Inc., n.d.). We tested for differences across populations and regions in the response variables of leaf length, panicle length, panicle number, percent recovery from desiccation, and genome size using generalized mixed linear models. Plant ID was included in each model as a random effect and population was nested within region. We tested for differences in morphological traits (leaf length, panicle length, panicle number) separately for field collected plants (parental generation) and plants cultivated in common conditions (progeny generation). To estimate heritability in leaf length, panicle length, and panicle number we computed the R2 value across field (parental) and common garden (progeny) plants by running separate mixed effects linear models to test the effect of parental phenotypes (field plants) on progeny phenotypes (common garden plants) with parental plant ID included in the model as a random effect. Desiccation responses were also quantified separately for field collected and common garden plants because different response variables were measured in the two desiccation assays. For field collected plants, we tested for differences in *F*_*v*_*/F*_*m*_, and RWC with generalized linear models that included the fixed effects of region, timepoint, and the interaction between the two. For plants cultivated under common conditions we tested for differences in *F*_*v*_*/F*_*m*_ and percent recovery. We tracked *F*_*v*_*/F*_*m*_ throughout the entire timecourse and tested for differences across regions using a repeated measures ANOVA. For percent recovery, we used a mixed effects linear model to test the effect of population and region on recovery at 192 hours post rehydration, with plant ID included as a random effect.

Dimension reduction via principal component analyses (PCA) was used to visualize emergent patterns in this multivariate dataset. Trait associations and correlations were computed by taking the mean value of leaf length, panicle length, panicle number, genome size, and percent recovery for each accession, and the site characteristics of elevation, annual precipitation, and annual temperature. A covariance matrix was generated by restricted maximum likelihood (REML) across all of these factors and singular value decomposition (SVD) was used to obtain eigenvectors and eigenvalues for PCA.

## RESULTS

### Field collections and study sites

We collected 33 accessions of *M. caffra* across 11 populations in four major areas of Mpumalanga and Limpopo provinces in South Africa. Study sites range from low elevation xeric to high elevation mesic sites and span ∼500 km in distance, 720 m in elevation, ∼400 mm in mean annual rainfall, and 3°C in mean annual temperature (Marks et al., 2022) (Figure 1; Supplementary Table S1). In general, sites within Buffelskloof are the wettest, coolest, and highest elevation. Sites within Veloren and Sister Mountains are intermediate, and sites within Swebe Swebe are the hottest, driest, and lowest elevation.

### Morphological and reproductive phenotypes parallel environmental gradients

Significant morphological and reproductive differences across study sites were identified in field specimens. In general, plants from higher elevation mesic areas were smaller with shorter leaves (p<0.0001) and panicles (p<0.0001) relative to plants from lower elevation xeric areas which were larger, with longer leaves and panicles (Figure 2). Interestingly, the smaller plants from higher elevation mesic sites produced more panicles than the larger plants from lower elevation xeric sites (p=0.0012).

**Figure 2.**
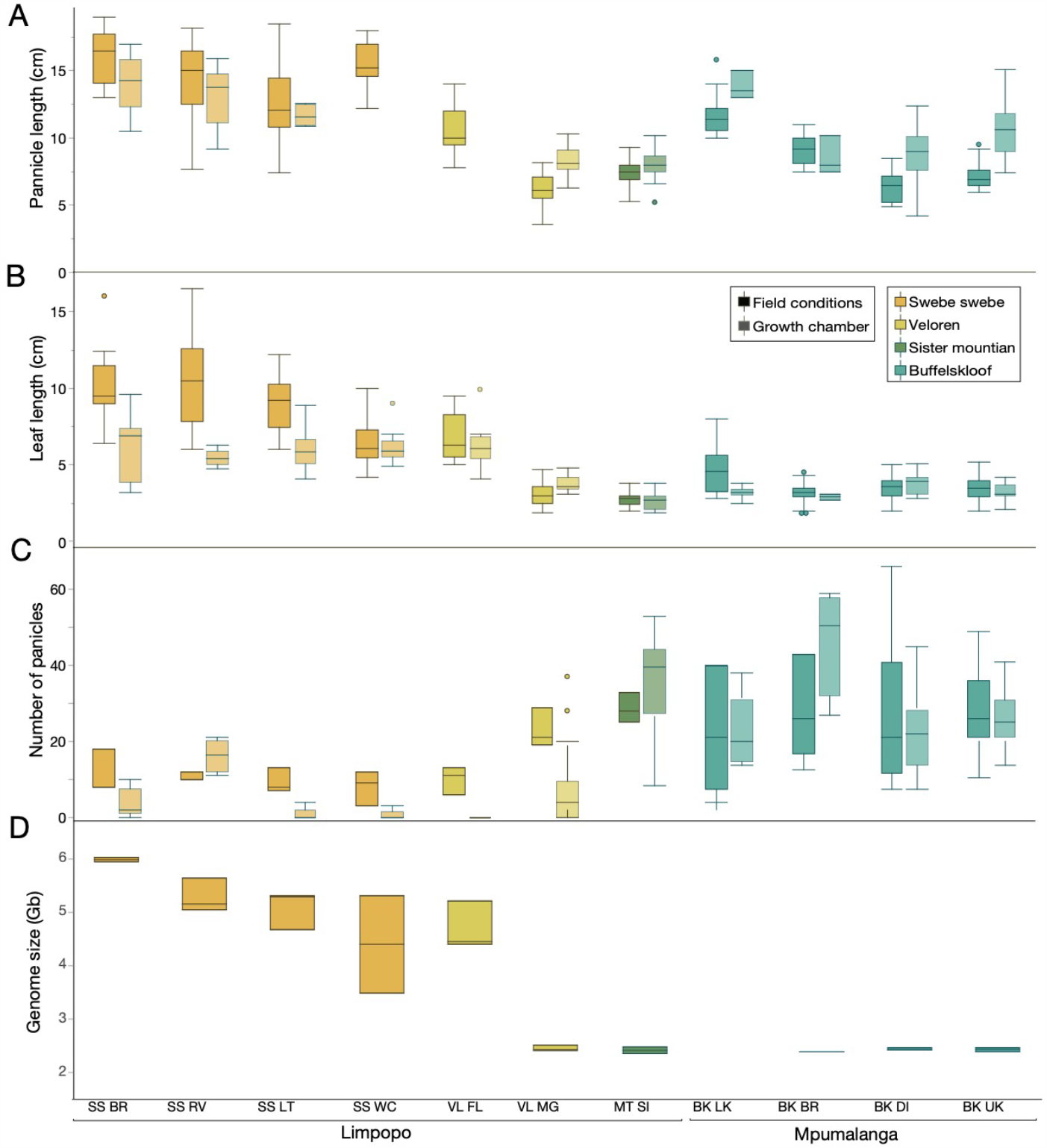
Morphological and reproductive phenotypes of plants in field and common garden conditions. Regions are ordered from xeric to mesic and sites are ordered from low to high elevation within each region. A) mean panicle number, B) mean panicle length, and C) mean leaf length are shown for both field and common garden plants. D) Genome size variation across the study area. SS=Swebe Swebe, VL=Veloren, MT SI=Mountain sister, and BK=Buffelskloof.

Phenotypic differences were persistent when plants were cultivated under common conditions. Paralleling field phenotypes, plants from the high elevation mesic sites were consistently smaller with more panicles than plants from lower elevation xeric sites which were larger, with longer leaves (p<0.0001) and panicles (p<0.0001), but fewer overall panicles (p<0.0001) (Figure 2A-C). Correlations between parental and progeny phenotypes were tight, suggesting a high degree of heritability in morphological traits. R2 values were 0.837 for leaf length, 0.662 for panicle length, and 0.859 for panicle number. These findings suggest a high degree of stability in morphological and reproductive phenotypes in these accessions. Taken together, these data support the existence of two diverging ecotypes that occur along an environmental gradient–a small reproductive ecotype from high elevation mesic sites, and a large but less reproductive ecotype from low elevation xeric sites.

An estimated 90% of chloridoid grasses are polyploid (Davidse et al., 1986; Roodt and Spies, 2003), and we tested if differences in ploidy existed across *M. caffra* populations and if they were associated with variation in phenotypic traits. Initially, we quantified the ploidy of a single representative accession from Buffelskloof using a K-mer approach with whole genome sequencing data. The K-mer plot shows six distinct peaks, indicating this accession is hexaploid (2n=6x), with a monoploid genome size of ∼300-400 Mb (Figure S1). This relatively small monoploid genome size is similar to other chloridoid grasses, which tend to have more compact genomes than most grass lineages (VanBuren et al., 2015, 2018, 2020; Pardo et al., 2020; Chávez Montes et al., 2022). The distribution of K-mers in the *M. caffra* genome and the prevalence of ‘aaab’ over ‘aabb’ alleles is consistent with autopolyploidy and high genetic diversity (Ranallo-Benavidez et al., 2020).

We then used flow cytometry to estimate genome size of each of the 33 *M. caffra* accessions included in the current study. We observed substantial variation in genome sizes across populations and regions (Figure 2D). Genome sizes varied by more than two-fold from 2.35 to 6.04 Gb per 2C, with the larger plants from lowland xeric sites having consistently and significantly larger genome sizes compared to the smaller plants from the high elevation mesic sites (p<0.0001). We estimate that ploidy varies from hexaploid (2n=6x) in the high elevation ecotype to dodecaploid (2n=12x) or more in the low elevation ecotype. Within the xeric low elevation ecotype, genome size ranges from 3.5 to 6 Gb per 2C so higher order ploidies above dodecaploid are certainly possible.

### Higher order polyploids exhibit enhanced recovery from desiccation

*M. caffra* plants from the field and their selfed progeny grown under common conditions were subjected to controlled drying in separate desiccation assays. Comprehensive desiccation assays were performed on a replicated set of all accessions grown from seed under common conditions. Mature seed grown plants were dried slowly over the course of 28 days and their ability to recover was assessed. The endpoint RWC of all samples was confirmed at less than 10%, indicating that all plants were fully desiccated prior to rehydration.

*F*_*v*_*/F*_*m*_ declined at similar rates and recovered to near baseline levels in all plants from all sites (Figure 3B). There were subtle differences in the rate of recovery after rehydration, but when integrated across the entire timecourse these differences were insignificant, suggesting that recovered tissues are functional in all populations. Because *F*_*v*_*/F*_*m*_ is a relatively insensitive measure, the subtle differences detected here may translate into more substantial differences in gas exchange and carbon assimilation (Seaton and Walker, 1990).

**Figure 3.**
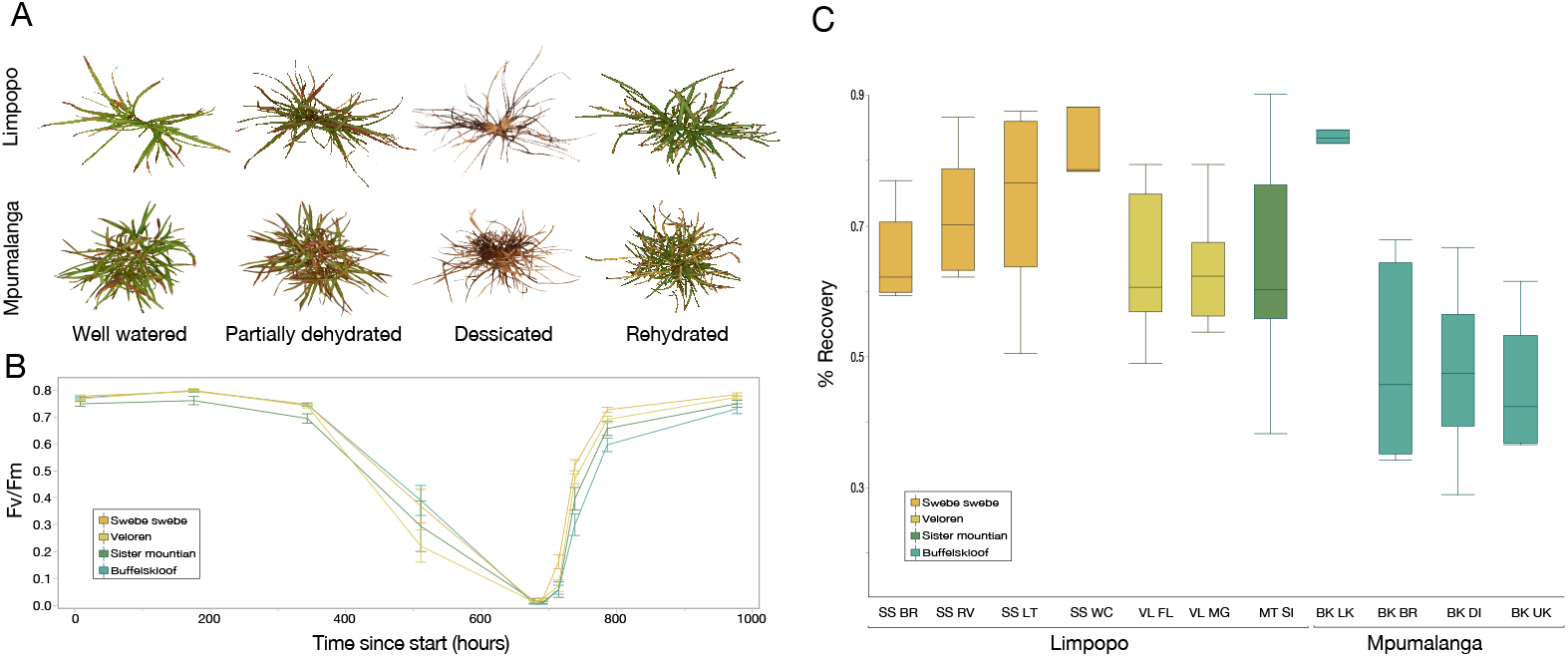
**A)** Images of representative plants throughout the desiccation and rehydration timecourse. **B)** Photochemical efficiency of Photosystem II (*F*_*v*_*/F*_*m*_) of plants during a controlled drying timecourse. **C)** The proportion of tissue that recovered after desiccation was variable across populations. Plants from the most xeric study sites recovered more completely from desiccation compared to plants from mesic sites. Regions are ordered from xeric to mesic and sites are ordered from low to high elevation within each region.

Despite the similar speed of drying and recovery, we detected significant differences across sites (p=0.0002) and regions (p=0.0004) in the proportion of tissue that recovered from desiccation (Figure 3C). In general, plants from high elevation mesic regions suffered more senescence and tissue loss than plants from lowland xeric sites (Figure 3A). Within regions, plants from more mesic areas exhibited more senescence than plants from xeric areas. For example, BK-LK is one of the driest sites within the generally mesic region of Buffelskloof, and those plants had higher recovery than morphologically similar plants from nearby sites. Because these data were generated from plants cultivated in common conditions, they suggest that genetic, rather than plastic, differences between populations drive differences in recovery from desiccation.

In field collected plants, the maximum quantum yield of Photosystem II (*F*_*v*_*/F*_*m*_) and leaf relative water content (RWC) dropped to a lower absolute level in plants from the lowland xeric sites (Swebe Swebe, Limpopo) relative to plants from the higher elevation mesic sites (Buffelskloof, Mpumalanga) (Figure S2). However, the lowland plants recovered more rapidly than plants from the high elevation mesic sites, represented by the significant interaction term of region and timepoint for both *F*_*v*_*/F*_*m*_ (P<0.0001) and RWC (P=0.0483) (Figure S2).

Histological differences in subcellular morphology were also evident between these two populations of *M. caffra*. In the more mesic Mpumalanga environment, high starch levels were visible in hydrated tissue whereas none were evident in plants from the more xeric Limpopo site (Figure 4A and 4C). In addition, evidence of a response to water deficit stress, or a memory of such a stress, is notable in that the vacuoles of the plants from Limpopo showed accumulation of electron dense compatible solutes in hydrated conditions, a proposed mechanism for stabilization against mechanical stress in resurrection plants (Farrant et al., 2007). Plants from Limpopo reached a desiccated state after 7 days of dehydration, whereas those from Mpumalanga had reached only ∼30% RWC at this stage. Subcellular organization of the former was well preserved, with little evidence of plasmalemma withdrawal, well defined plasmodesmata, intact organelles, and a densely stained cytoplasm (Figure 4D), typical of the desiccated, quiescent state reported for other resurrection angiosperms (Farrant et al., 2007). Survival below 40% RWC is considered the defining feature of desiccation tolerance (Giarola et al., 2017) and thus although the plants from Mpumalanga did not reach an air-dry state in this experiment, it is clear from ultrastructural studies that tissues are likely viable, but yet to undergo the massive re-organization and stabilization of cytoplasm that occurs in resurrection plants during the loss of this final ∼35% water. While there is some evidence of plasmalemma withdrawal in drying tissues from Mpumalanga, this is likely due to the mildly aqueous fixation protocol used. There is no evidence of plasmalemma damage, and all organelles were intact. Chloroplasts in drying leaves from Mpumalanga resembled those in the desiccated leaves of the Limpopo population, and importantly, the mitochondria appear in a highly active state (Figure 4B), typical of entering the final stages of desiccation in resurrection plants (Farrant et al., 2017).

**Figure 4.**
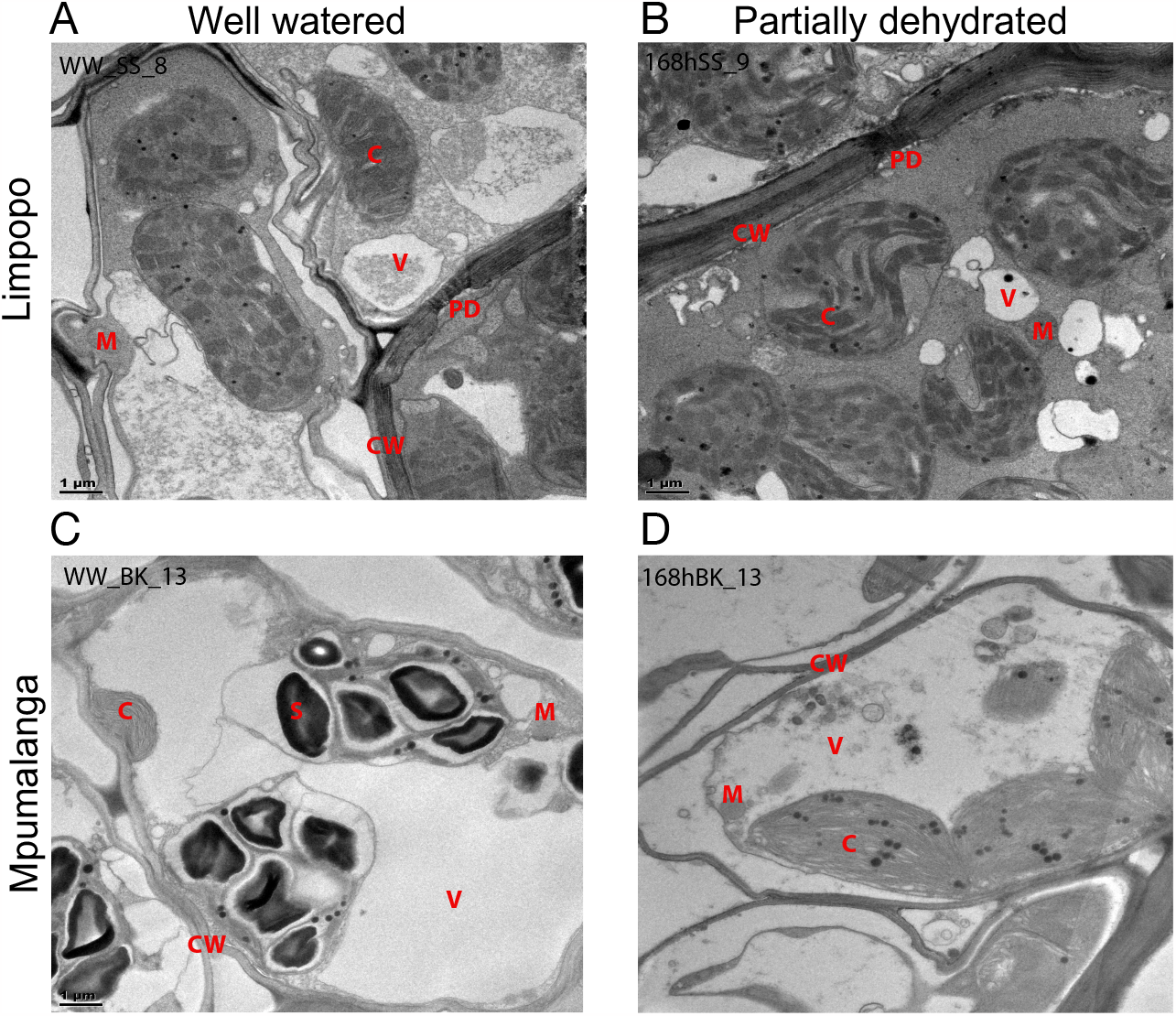
Transmission electron micrographs of mesophyll cells of plants collected from Mpumalanga (C and D) and Limpopo (A and B) under fully hydrated conditions (A and C) and desiccated or imminent desiccation (30% RWC) of these populations (B and D). Abbreviations: CW=cell wall; C=chloroplast; M=mitochondrion; PD=plasmodesmata; S=starch; V=vacuole

**Figure 5.**
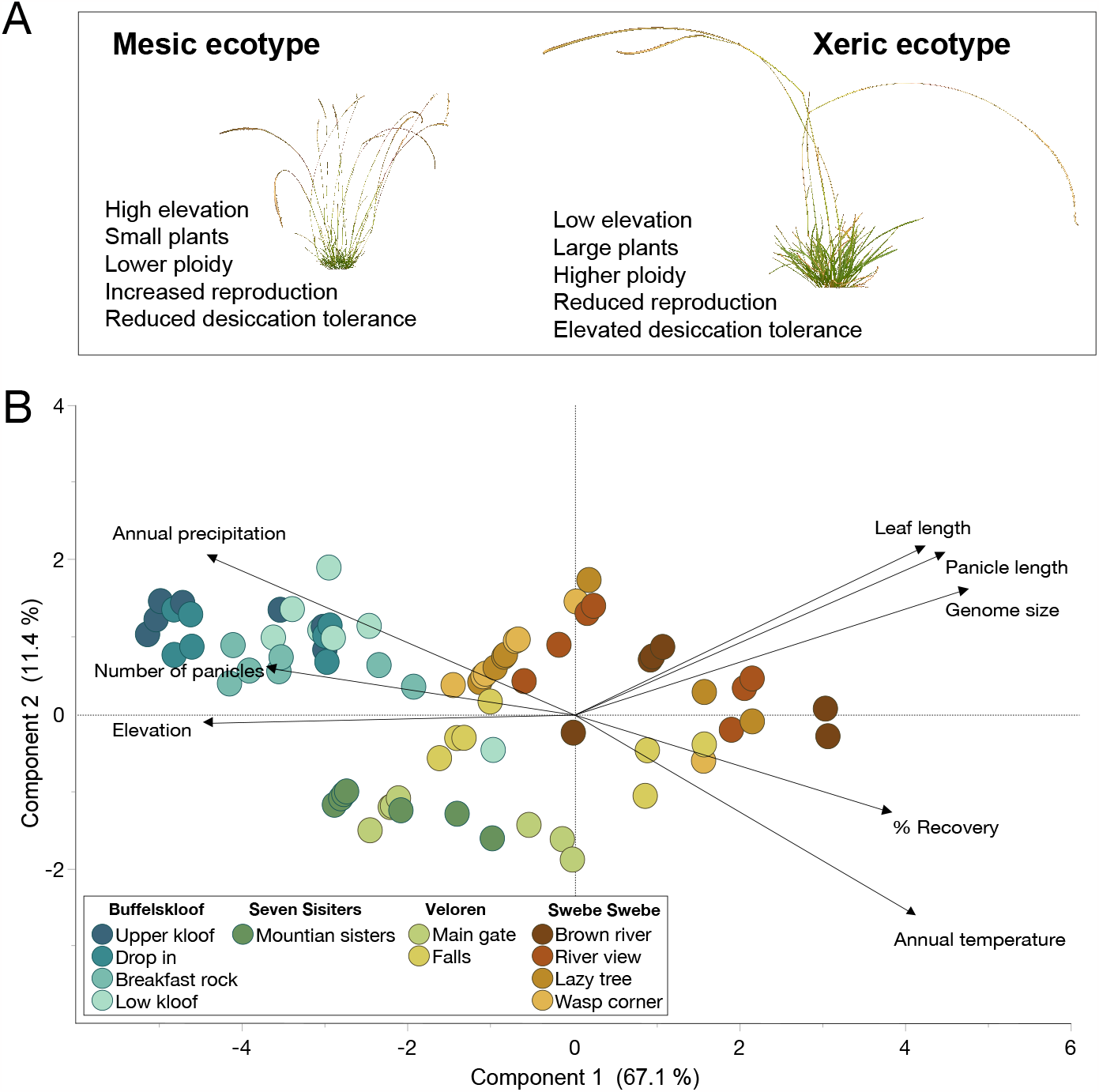
**A)** Summary of ecotype differences across the study area. **B)** Principal component analysis (PCA) of leaf length, panicle length, panicle number, recovery from desiccation, genome size, elevation, annual precipitation, and annual temperature. Samples are colored by site.

### Functional syndromes delineating the mesic and xeric *M. caffra* ecotypes

We identified numerous positively and negatively correlated traits. Taken together, we describe two ecotypes of *M. caffra* that occur along an environmental gradient in water availability and elevation. Plants from high elevation mesic sites are generally small, highly reproductive, with reduced desiccation tolerance, and lower ploidy. In contrast, plants from the xeric lowland sites are larger, less reproductive, more desiccation tolerant, and have higher ploidy. Leaf length, panicle length, and genome size were strongly positively correlated. Percent recovery was positively associated with genome size (R=0.52), panicle length (R=0.58) and leaf length (R=0.46), but negatively associated with number of panicles (R=-0.54) and elevation (R=-0.57). Elevation was positively associated with precipitation (R=-0.63) and number of panicles (R=0.54), but negatively correlated with temperature (R=-0.70), genome size (R=-0.74), panicle length (R=-0.68) and leaf length (R=-0.70) (Table 1). PCA explained much of the variation in the dataset with PC1 accounting for 67.1% of the variability in the data and separating samples along a gradient of elevation, moisture, genome size, and desiccation tolerance. PC2 explains another 11.4% of the variability in these data and separated samples on more subtle differences in morphology.

**Table 1.** Environmental and trait correlations were generated by restricted maximum likelihood (REML) and used for Principal component analysis (PCA). Positive correlations are shown in red and negative correlations are shown in blue.

|  | *Annual precip.* | *Annual temp.* | *Elevation* | *Recovery* | *Genome size* | *Leaf length* | *Panicle length* | *Number of panicles* |
| --- | --- | --- | --- | --- | --- | --- | --- | --- |
| *Annual precip.* | 1.00 |  |  |  |  |  |  |  |
| *Annual temp.* | -0.90 | 1.00 |  |  |  |  |  |  |
| *Elevation* | 0.63 | -0.70 | 1.00 |  |  |  |  |  |
| *Recovery* | -0.61 | 0.57 | -0.57 | 1.00 |  |  |  |  |
| *Genome size* | -0.70 | 0.58 | -0.74 | 0.52 | 1.00 |  |  |  |
| *Leaf length* | -0.61 | 0.48 | -0.70 | 0.46 | 0.92 | 1.00 |  |  |
| *Panicle length* | -0.52 | 0.46 | -0.68 | 0.58 | 0.83 | 0.77 | 1.00 |  |
| *Number of panicles* | 0.58 | -0.45 | 0.54 | -0.54 | -0.60 | -0.58 | -0.39 | 1.00 |

## DISCUSSION

Polyploidy has enabled the evolution of complex and emergent traits (Huminiecki and Conant, 2012), including many adaptations related to diversification and resilience in plants (Otto, 2007; Te Beest et al., 2012; Soltis et al., 2015). Here, we tested the hypothesis that polyploidy enabled the evolution of desiccation tolerance (Giarola et al., 2017) using a plant system exhibiting natural variation along an environmental gradient. We identified heritable variation in desiccation tolerance in the resurrection grass *Microchloa caffra* that is associated with differences in both ploidy and environmental conditions. Accessions with higher ploidy were found at drier sites and displayed greater desiccation tolerance compared to their lower-ploidy counterparts. Polyploidy could enhance desiccation tolerance through multiple mechanisms. For instance, duplicated genes can neo-functionalize to acquire novel functions (Lynch and Conery, 2000) that may enhance desiccation tolerance. Polyploidy can also increase gene dosage (Adams and Wendel, 2005), leading to higher abundance of enzymes, structural proteins, and critical end-point metabolites that play important roles in desiccation tolerance. Our data provide intriguing evidence that desiccation tolerance may be mediated by changes in ploidy and highlight the significance of polyploidy as a potential driver of the evolution of complex traits.

Broadly, we describe two distinct ecotypes of *M. caffra* that exhibit consistent differences in morphological, reproductive, and desiccation tolerance traits in both field and common growth conditions. Plant phenotypes varied along the environmental gradient of elevation, temperature, and water availability as predicted. We observed elevated desiccation tolerance and improved recovery outcomes in plants from the driest sites, compared to plants from more mesic sites. These data hint at local adaptation and suggest that strong selective pressures in arid environments favor the evolution of enhanced desiccation tolerance. In contrast, plants from mesic sites displayed reduced desiccation tolerance, a possible consequence of relaxed selective pressure on desiccation tolerance in these wetter environments. Similar patterns have been documented in other plant species (Marks et al., 2016, 2019a; Wang et al., 2018) supporting the notion that adaptation to different moisture regimes can lead to contrasting tolerance levels (Gutschick and BassiriRad, 2003). The ranges of these two ecotypes are not fully quantified and parallel assessments within the range of each ecotype could provide additional insight into whether environmental differences or increased ploidy are more important drivers of local adaptation in *M. caffra*.

Differences in reproductive allocation across the study area trended in the opposite direction of desiccation tolerance traits, with more reproductive plants occurring in the most mesic sites. Variation in ploidy levels may contribute to the observed reproductive differences, as polyploidy can interfere with meiotic processes that are essential for sexual reproduction and thereby result in reduced panicle and seed number (Comai, 2005; Herben et al., 2017), as observed here. In addition to the effects of polyploidy, reproductive differences across populations could be driven by selection against seed production in xeric sites due to frequent droughts that interrupt development. Alternatively, there may also be a tradeoff between desiccation tolerance and reproduction as has been suggested in the liverwort *Marchantia inflexa* (Marks et al., 2019a; b, 2020). The differences in reproduction observed here, parallel findings in the eudicot resurrection plant *Myrothamnus flabellifolia*, which was sampled along the same environmental gradient (Marks et al., 2022). However, in *M. flabellifolia*, desiccation tolerance is maintained at high levels at all sites (Marks et al., 2022), suggesting that tradeoffs between desiccation tolerance and reproduction are not universal in resurrection plants.

The role of polyploidy in the evolution of desiccation tolerance in plants is an open question, and although we provide evidence that polyploidy enhances tolerance in *M. caffra*, there are important confounding factors to consider. Most Chloridoid grasses (∼90%) are polyploid, making it difficult to distinguish causal and correlative relationships when placing ploidy within the context of emergent traits. Desiccation tolerant Chloridoids within *Oropetium* and *Tripogon* are generally diploid with small, compact genomes, suggesting polyploidy is not necessary for evolving desiccation tolerance within this clade. More broadly, natural polyploids arise mostly through unreduced gametes (Soltis and Soltis, 2009), and environmental stresses increase the rate of these errors during meiosis (Mason et al., 2011; Pecrix et al., 2011; De Storme et al., 2012). The extreme environments that resurrection plants inhabit and the repeated cycles of drying may increase the rate of polyploid formation within these species, rather than enabling this trait (Gutschick and BassiriRad, 2003). Instead of a declarative link, we suggest a more nuanced role of polyploidy where genome duplication improves desiccation tolerance traits by providing more raw genetic material for possible sub- and neo-functionalization and/or increasing the abundance of critical end-point metabolites, enzymes, and structural proteins (Giarola et al., 2017). Simply being polyploid is unlikely related to the evolution of desiccation tolerance, but our data suggest that it could be a mechanism for enhancing existing tolerance mechanisms and driving local adaptation. These complex interactions illustrate how evolutionary processes associated with genome duplication might shape a plant’s ability to thrive under abiotic stress. Further investigations of specific molecular mechanisms underlying these phenomena will provide insight into the interplay between polyploidy, desiccation tolerance, and adaptation to water scarcity. We suggest that *M. caffra* is a promising system for exploring the role of polyploidy in facilitating the rapid evolution of desiccation tolerance.

## ACKNOWLEDGEMENTS

This work was funded by NSF IOS grant no. PRFB-1906094 to RAM and DBI grant no. 2213983 to the Water and Life Interface Institute. We thank landowners and stewards Pieter and Jennie Pretorius, Ken Maude, Pieter Vervoort, Syd Catton, and Felicite Jackson for access to study sites, and Wayne and Messiah Mudenda for assistance with field logistics. We thank the South African National Herbarium, Pretoria for assistance with identification and vouchering of specimens.

## AUTHOR CONTRIBUTIONS

RAM, RV and JMF conceived of the study. RAM, PD, GMM, and KC contributed to data acquisition and curation. RAM conducted data analyses. RAM, PD, GMM, RV, and JMF contributed to data interpretation and conceptual framing of the manuscript. RAM drew the figures and wrote the manuscript. All authors edited and reviewed the manuscript.

## DATA AVAILABILITY

Data associated with this study are provided as a supplementary appendix.

## REFERENCES

Adams, K. L., and J. F. Wendel. 2005. Polyploidy and genome evolution in plants. Current opinion in plant biology 8: 135–141.

Alpert, P. 2006. Constraints of tolerance: why are desiccation-tolerant organisms so small or rare? The Journal of experimental biology 209: 1575–1584.

Bartels, D., and M. Z. M. Mattar. 2002. Oropetium thomaeum. A resurrection grass with a diploid genome. Maydica.

Bewley, J. D. 1979. Physiological Aspects of Desiccation Tolerance. Annual review of plant physiology 30: 195–238.

Bewley, J. D., and J. E. Krochko. 1982. Desiccation-Tolerance. In O. L. Lange, P. S. Nobel, C. aB. Osmond, and H. Ziegler [eds.], Physiological Plant Ecology II: Water Relations and Carbon Assimilation, 325–378. Springer Berlin Heidelberg, Berlin, Heidelberg.

Chávez Montes, R. A., A. Haber, J. Pardo, R. F. Powell, U. K. Divisetty, A. T. Silva, T. Hernández-Hernández, et al. 2022. A comparative genomics examination of desiccation tolerance and sensitivity in two sister grass species. Proceedings of the National Academy of Sciences of the United States of America 119.

Comai, L. 2005. The advantages and disadvantages of being polyploid. Nature reviews. Genetics 6: 836–846.

Cooper, K., and J. M. Farrant. 2002. Recovery of the resurrection plant Craterostigma wilmsii from desiccation: protection versus repair. Journal of experimental botany 53: 1805–1813.

Costa, M.-C. D., M. A. S. Artur, J. Maia, E. Jonkheer, M. F. L. Derks, H. Nijveen, B. Williams, et al. 2017. A footprint of desiccation tolerance in the genome of Xerophyta viscosa. Nature Plants 3: 17038.

Davidse, G., T. Hoshino, and B. K. Simon. 1986. Chromosome counts of Zimbabwean grasses (Poaceae) and an analysis of polyploidy in the grass flora of Zimbabwe. South African journal of botany: official journal of the South African Association of Botanists = Suid-Afrikaanse tydskrif vir plantkunde: amptelike tydskrif van die Suid-Afrikaanse Genootskap van Plantkundiges 52: 521–528.

De Storme, N., G. P. Copenhaver, and D. Geelen. 2012. Production of diploid male gametes in Arabidopsis by cold-induced destabilization of postmeiotic radial microtubule arrays. Plant physiology 160: 1808–1826.

Farrant, J. M., W. Brandt, and G. G. Lindsey. 2007. An Overview of Mechanisms of Desiccation Tolerance in Selected Angiosperm Resurrection Plants. Plant Stress 1: 72–84.

Farrant, J.M., Cooper, K., Dace, H.J.W.S, Bentely, J., Hilgart, A. 2017. Desiccation tolerance. In S. Shabala [ed.], Plant Stress Physiology, 217–252. CABI, Wallingford, UK.

Fick, S. E., and R. J. Hijmans. 2017. WorldClim 2: new 1-km spatial resolution climate surfaces for global land areas. International Journal of Climatology 37: 4302–4315.

Giarola, V., Q. Hou, and D. Bartels. 2017. Angiosperm Plant Desiccation Tolerance: Hints from Transcriptomics and Genome Sequencing. Trends in plant science 22: 705–717.

Gutschick, V. P., and H. BassiriRad. 2003. Extreme events as shaping physiology, ecology, and evolution of plants: toward a unified definition and evaluation of their consequences. The New phytologist 160: 21–42.

Herben, T., J. Suda, and J. Klimešová. 2017. Polyploid species rely on vegetative reproduction more than diploids: a re-examination of the old hypothesis. Annals of botany 120: 341–349.

Huminiecki, L., and G. C. Conant. 2012. Polyploidy and the evolution of complex traits. International journal of evolutionary biology 2012: 292068.

Lynch, M., and J. S. Conery. 2000. The evolutionary fate and consequences of duplicate genes. Science 290: 1151–1155.

Marçais, G., and C. Kingsford. 2011. A fast, lock-free approach for efficient parallel counting of occurrences of k-mers. Bioinformatics 27: 764–770.

Marks, R. A., J. F. Burton, and D. N. McLetchie. 2016. Sex differences and plasticity in dehydration tolerance: insight from a tropical liverwort. Annals of botany 118: 347–356.

Marks, R. A., J. M. Farrant, D. Nicholas McLetchie, and R. VanBuren. 2021. Unexplored dimensions of variability in vegetative desiccation tolerance. American journal of botany.

Marks, R. A., M. Mbobe, M. Greyling, J. Pretorius, D. N. McLetchie, R. VanBuren, and J. M. Farrant. 2022. Variability in Functional Traits along an Environmental Gradient in the South African Resurrection Plant Myrothamnus flabellifolia. Plants 11.

Marks, R. A., B. D. Pike, and D. Nicholas McLetchie. 2019a. Water stress tolerance tracks environmental exposure and exhibits a fluctuating sexual dimorphism in a tropical liverwort. Oecologia 191: 791–802.

Marks, R. A., J. J. Smith, Q. Cronk, C. J. Grassa, and D. N. McLetchie. 2019b. Genome of the tropical plant Marchantia inflexa: implications for sex chromosome evolution and dehydration tolerance. Scientific reports 9: 8722.

Marks, R. A., J. J. Smith, R. VanBuren, and D. N. McLetchie. 2020. Expression dynamics of dehydration tolerance in the tropical plant Marchantia inflexa. The Plant journal: for cell and molecular biology.

Mason, A. S., M. N. Nelson, G. Yan, and W. A. Cowling. 2011. Production of viable male unreduced gametes in Brassica interspecific hybrids is genotype specific and stimulated by cold temperatures. BMC plant biology 11: 103.

Mehra, P. N., and M. L. Sharma. 1975. Cytological Studies in Some Central and Eastern Himalayan Grasses. Cytologia 40: 453–462.

Oliver, M. J., J. M. Farrant, H. W. M. Hilhorst, S. Mundree, B. Williams, and J. D. Bewley. 2020. Desiccation Tolerance: Avoiding Cellular Damage During Drying and Rehydration. Annual review of plant biology.

Oliver, M. J., Z. Tuba, and B. D. Mishler. 2000. The evolution of vegetative desiccation tolerance in land plants. Plant Ecology 151: 85–100.

Oliver, M. J., J. Velten, and B. D. Mishler. 2005. Desiccation tolerance in bryophytes: a reflection of the primitive strategy for plant survival in dehydrating habitats? Integrative and comparative biology 45: 788–799.

Otto, S. P. 2007. The evolutionary consequences of polyploidy. Cell 131: 452–462.

Pardo, J., C. Man Wai, H. Chay, C. F. Madden, H. W. M. Hilhorst, J. M. Farrant, and R. VanBuren. 2020. Intertwined signatures of desiccation and drought tolerance in grasses. Proceedings of the National Academy of Sciences of the United States of America 117: 10079–10088.

Pecrix, Y., G. Rallo, H. Folzer, M. Cigna, S. Gudin, and M. Le Bris. 2011. Polyploidization mechanisms: temperature environment can induce diploid gamete formation in Rosa sp. Journal of experimental botany 62: 3587–3597.

Porembski, S. 2011. Evolution, Diversity, and Habitats of Poikilohydrous Vascular Plants. In D. Bartels, U. Lüttge, and E. Beck [eds.], Plant Desiccation Tolerance, 139–156.

Proctor, M. C. F., and Z. Tuba. 2002. Poikilohydry and homoihydry: antithesis or spectrum of possibilities? The New phytologist 156: 327–349.

Ranallo-Benavidez, T. R., K. S. Jaron, and M. C. Schatz. 2020. GenomeScope 2.0 and Smudgeplot for reference-free profiling of polyploid genomes. Nature communications 11: 1432.

Roodt, R., and J. J. Spies. 2003. Chromosome studies in the grass subfamily Chloridoideae. I. Basic chromosome numbers. TAXON 52: 557–583.

SAS Institute Inc. JMP®, Version 10. Cary, NC.

Seaton, G. G. R., and D. A. Walker. 1990. Chlorophyll Fluorescence as a Measure of Photosynthetic Carbon Assimilation. Proceedings. Biological sciences / The Royal Society 242: 29–35.

Soltis, D. E., C. J. Visger, and P. S. Soltis. 2014. The polyploidy revolution then舰and now: Stebbins revisited. American journal of botany 101: 1057–1078.

Soltis, P. S., D. B. Marchant, Y. Van de Peer, and D. E. Soltis. 2015. Polyploidy and genome evolution in plants. Current opinion in genetics & development 35: 119–125.

Soltis, P. S., and D. E. Soltis. 2009. The role of hybridization in plant speciation. Annual review of plant biology 60: 561–588.

Te Beest, M., J. J. Le Roux, D. M. Richardson, A. K. Brysting, J. Suda, M. Kubešová, and P. Pyšek. 2012. The more the better? The role of polyploidy in facilitating plant invasions. Annals of botany 109: 19–45.

Tebele, S. M., R. A. Marks, and J. M. Farrant. 2021. Two Decades of Desiccation Biology: A Systematic Review of the Best Studied Angiosperm Resurrection Plants. Plants 10.

VanBuren, R. 2017. Desiccation tolerance: Seedy origins of resurrection. Nature Plants 3: 17046.

VanBuren, R., D. Bryant, P. P. Edger, H. Tang, D. Burgess, D. Challabathula, K. Spittle, et al. 2015. Single-molecule sequencing of the desiccation-tolerant grass Oropetium thomaeum. Nature 527: 508–511.

VanBuren, R., C. Man Wai, X. Wang, J. Pardo, A. E. Yocca, H. Wang, S. R. Chaluvadi, et al. 2020. Exceptional subgenome stability and functional divergence in the allotetraploid Ethiopian cereal teff. Nature communications 11: 884.

VanBuren, R., C. M. Wai, J. Keilwagen, and J. Pardo. 2018. A chromosome-scale assembly of the model desiccation tolerant grass Oropetium thomaeum. Plant Direct 2: e00096.

VanBuren, R., C. M. Wai, Q. Zhang, and X. Song. 2017. Seed desiccation mechanisms coopted for vegetative desiccation in the resurrection grass Oropetium thomaeum. Plant, cell &.

Wang, X., Z.-H. Chen, C. Yang, X. Zhang, G. Jin, G. Chen, Y. Wang, et al. 2018. Genomic adaptation to drought in wild barley is driven by edaphic natural selection at the Tabigha Evolution Slope. Proceedings of the National Academy of Sciences of the United States of America 115: 5223–5228.

Wendel, J. F. 2000. Genome evolution in polyploids. Plant molecular biology 42: 225–249.

