## Supplementary table for "Polyploidy enhances desiccation tolerance in the grass *Microchloa caffra*"

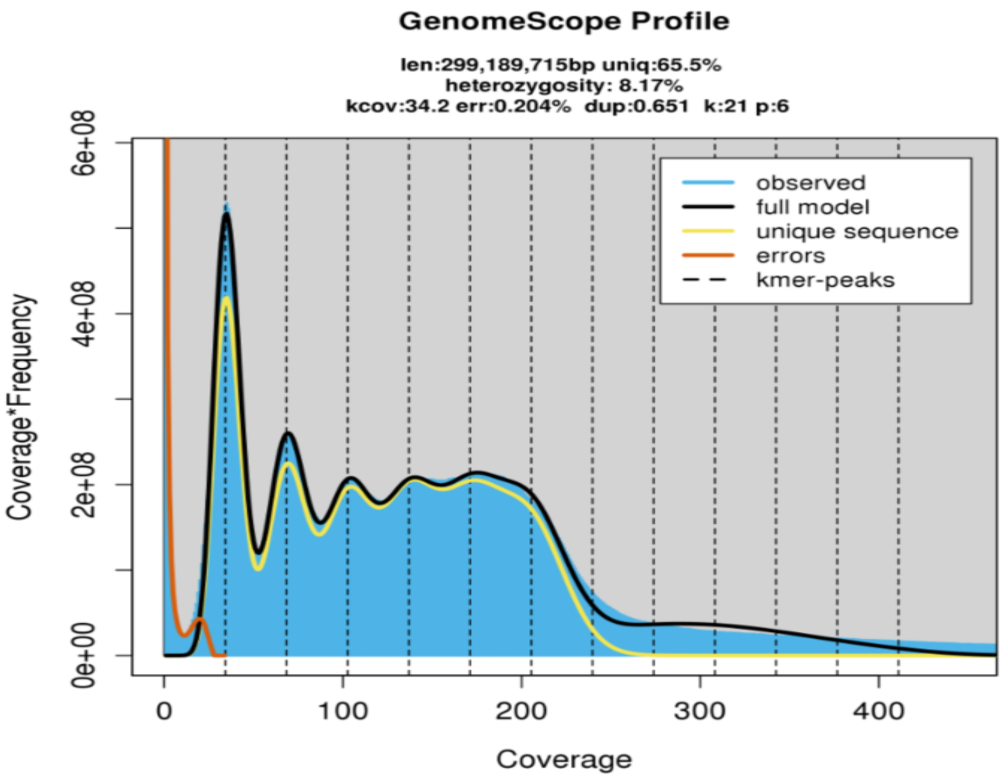


**Figure S1.** K-mer analyses of whole genome sequencing data.

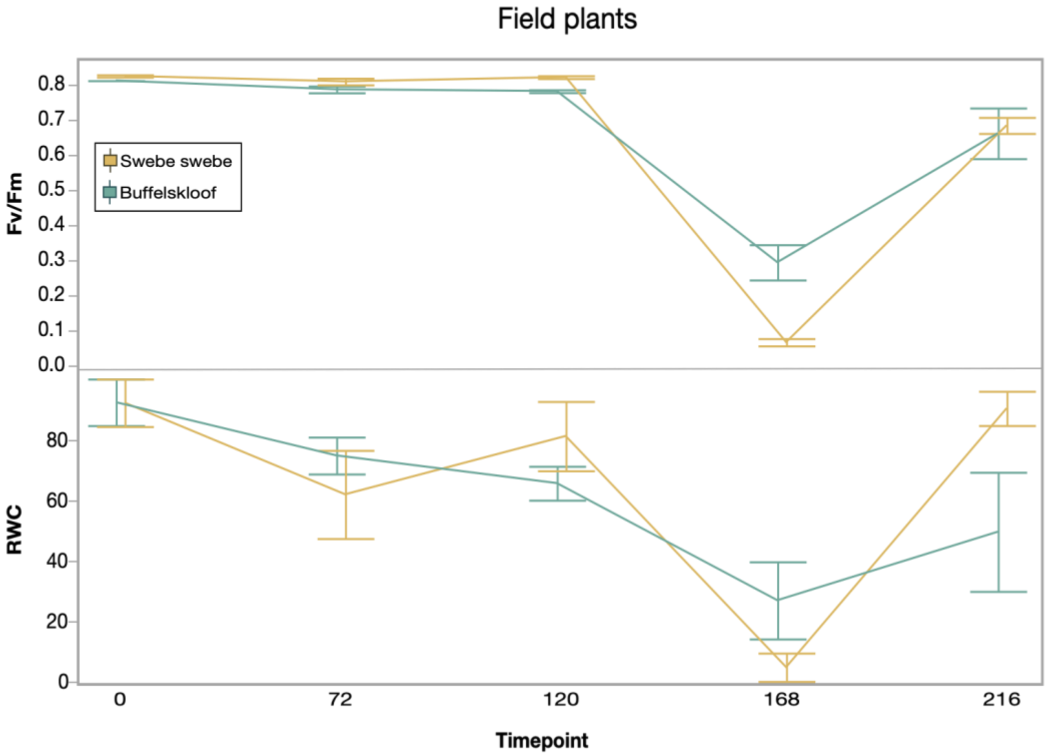


**Figure S2.** Changes in photochemical efficiency of PSII ($F_v/F_m$) and RWC of field collected plants during a desiccation treatment imposed at the University of Cape Town. Selected plants from two sites (5 from Buffelskloof and 6 from Swebe Swebe) were desiccated under controlled conditions and rehydrated after 168 hours. Error bars represent standard error of the mean.

**Table S1:** Collection locations, mean annual precipitation and temperature, geo coordinates, and elevation. Precipitation and temperature data were taken from worldclim (Fick and Hijmans, 2017). . SS=Swebe Swebe, VL=Veloren,, and BK=Buffelskloof.

| Population name | Annual precip (mm) | Annual temp (C) | GPS Coordinates | Elevation (M) |
|---|---|---|---|---|
| BK breakfast rock | 924 | 17 | 25.19.785 S, 030.29.650 E | 1304 |
| BK drop in | 939 | 15 | 25.18.041 S, 030.30.463 E, | 1515 |
| BK low kloof | 904 | 17 | 25.20.085 S, 030.29.385 E | 1193 |
| BK upper kloof | 939 | 15 | 25.17.078 S, 030.30.579 E | 1631 |
| Mountain sister | 645 | 18.5 | 24.29.520 S, 027.44.377 E | 1498 |
| SS brown river | 513 | 19.7 | 23.45.032 S, 028.05.557 E | 911 |
| SS lazy tree | 627 | 17.8 | 23.48.731 S, 028.04.277 E | 1085 |
| SS river view | 580 | 18.4 | 23.47.418 S, 028.04.553 E | 1083 |
| SS wasp corner | 599 | 18.2 | 23.51.36 S, 028.02.188 E | 1278 |
| VL falls | 613 | 19 | 24.48.235 S, 028.22.622 E | 1295 |
| VL main gate | 623 | 18.7 | 24.47.624 S, 028.21.328 E | 1213 |
